# Assessment of distant-site rescue elements for CRISPR toxin-antidote gene drives

**DOI:** 10.1101/2023.01.06.522951

**Authors:** Jingheng Chen, Xuejiao Xu, Jackson Champer

**Affiliations:** Center for Bioinformatics, School of Life Sciences, Peking-Tsinghua Center for Life Sciences, Peking University, Beijing, China 100871

## Abstract

New types of gene drives promise to provide increased flexibility, offering many options for confined modification or suppression of target populations. Among the most promising are CRISPR toxin-antidote gene drives, which disrupt essential wild-type genes by targeting them with Cas9/gRNA, resulting in their removal. This increases the frequency of the drive in the population. All these drives, plus homing modification rescue drives, rely on having an effective rescue element, which consists of a recoded version of the target gene. This rescue element can be at the same site as the target gene, which maximizes the chance of efficient rescue, or at a distant site, which allows some other useful options, such as easily disrupting another essential gene or achieving greater confinement. Previously, we developed a homing rescue drive targeting a haplolethal gene and a toxin-antidote drive targeting an essential but haplosufficient gene. These successful drives had functional rescue elements but suboptimal drive efficiency. Here, we attempted to construct new toxin-antidote drives targeting these genes with a distant-site configuration from three different loci. We found that use of additional gRNAs increased cut rates to nearly 100%. However, all distant-site rescue elements failed for both haplolethal and haplosufficient target genes. Furthermore, one rescue element with a minimally recoded rescue element was used as a template for homology-directed repair for the target gene on a different chromosomal arm, resulting in the formation of functional resistance alleles at high frequency. Together, these results can inform the design of future CRISPR-based toxin-antidote gene drives.

## Introduction

CRISPR homing gene drives have been shown to rapidly spread through laboratory populations for purposes of population suppression^1^ or modification^2^. Such gene drives could be particularly valuable for the prevention of vector-borne disease, but they also have other applications such as the removal of invasive species and agricultural pests^3–5^. Yet, the best studied homing gene drives are “unconfined” in that a small release could lead to spread of the drive throughout the entire species, or at least all connected populations. This may be undesirable and could prevent deployment of gene drives when only specific populations should be targeted, such as removal of invasive species outside their native range^6–8^.

Several options exist for confined gene drives^3,5^, including *Medea*^9^, haploinsufficient underdominance^10^, chromosomal rearrangements^11,12^, incompatibility underdominance^13^, and *Wolbachia* cytoplasmic incompatibility drive^14^. Some of the latest and most promising confined drive designs are based on CRISPR nucleases and can avoid functional resistance alleles by use of multiplexed gRNAs. They work by targeting and disrupting an essential gene with Cas9 while also providing a recoded rescue copy of the target gene. Even though the gene drive is not copied by drive conversion as in homing drives, it still can increase in relative frequency by disrupting and removing wild-type alleles. These include the highly similar TARE (Toxin-Antidote Recessive Embryo)^15,16^ and ClvR (Cleave and Rescue)^17,18^ systems that are experimentally demonstrated and target a haplosufficient gene (in which one functioning copy is sufficient for an organism to retain high fitness). Such drives only gain an introduction threshold when they have fitness costs^19^, which refers to the necessary number of drive individuals that must be released for the drive to be successful. Introduction threshold usually serves as a good measure of how confined a drive is. Other forms of TARE drive that could likely be readily developed could provide for higher introduction thresholds, enabling flexible confinement options^20^ or options for self-limiting drive^21^ (similar to killer-rescue systems^22^). Together, these drives could serve as confined modification drive systems. They could also support confined suppression as part of a tethered drive system^16,23^ if sufficiently efficient homing suppression drives can be constructed in the target organism or even confined suppression alone if an adequate cargo could be developed^24^. Because development of these methods would be nontrivial, even if functional resistance alleles can be avoided, another option is to use a CRISPR toxin-antidote drive targeting a haplolethal gene (in which two functioning copies are required for an organism to have high fitness). Such TADE (Toxin-Antidote Dominant Embryo) drives could be used to provide a flexible level of confinement for both modification and suppression^19,20,25–27^. While TADE drives have not yet been constructed, a homing modification rescue drive targeting a haplolethal gene was successful in flies^2^.

One common need for all of these CRISPR toxin-antidote designs and also homing modification rescue drives, is an effective rescue element. This means that the gene is essentially equivalent to the wild-type version in terms of expression timing and amount, or at least close enough to avoid any significant fitness costs. There are several possible variations in any recoded region that could affect the chance of successful rescue. The first is the genomic location of the rescue element. In “same-site” rescue drives such as the experimental demonstration of TARE^15,16^ and most homing-based designs^2,28–30^, the recoded copy is placed at the same genomic locus as the original gene. This means that all upstream regulatory elements should be functional normally. In “distant-site” drives, the entire rescue element is not located at the same site as the wild-type genes. Though such rescue elements have also proven successful in ClvR systems^17,18^, some upstream regulatory elements may be missed if the synthetic promoter element is too small, and genomic structure at the new site could also affect expression. Another potential level of variation is the degree of recoding. A minimum, the gRNA sites themselves need to be recoded. Going further, adjacent coding areas could be recoded, and introns can be eliminated. Even further may involve changes to the 5’ or 3’ UTR (such as using a 3’UTR from a closely related species or a synthetic 3’UTR). More changes mean that potentially important specific sequences may be lost, changing the rescue element’s expression pattern, and reducing the chance of successful rescue. For homing drives and other same-site systems, recoding generally needs to encompass a large area on at least one side of the gRNA target sites to avoid undesired homology-directed repair of only the recoded region. In general, it is thought that for haplolethal genes that are more sensitive to level of expression, it may be more difficult to generate a successful rescue element.

Though efficient distant-site rescue elements may theoretically be more difficult to generate, distant-site drives have their advantages as well. Such drives can operate at a higher efficiency when Cas9 cleavage rates are very low^19^. More importantly, such drives can be placed in a second target gene (in which rescue is not desired), allowing the drive to disrupt the target gene without the need for additional gRNAs, simplifying drive construction for promising candidates such as TADE suppression drive^19,20,25^. It can also open up the possibility of targeting genes that would otherwise be worse candidates due to closer spacing between target sites and the drive insertion site (if the spacing is too close, then undesired homology-directed repair could form a functional resistance allele)^15^. Finally, it might be required for certain types of highly confined systems in which two rescue elements for different genes must be placed at the same genomic site (thus allowing at most one of these to be a same-site drive allele)^20^.

In this study, we had several objectives. We tested distant-site rescue in a variety of toxin-antidote systems, but we found that it was inefficient for both haplolethal and haplosufficient gene targets. However, we were able to greatly improve Cas9 cut rates of previous drive systems^2,15^ via use of additional gRNA target sites, despite constant Cas9 expression. When testing a minimally recoded rescue element, we found that undesired homology-directed repair could form functional resistance alleles based on this template, despite it being located on a different chromosomal arm as the drive target site. Together, these observations provide important lessons for the design of CRISPR toxin-antidote gene drives.

## Methods

### Plasmid construction

For plasmid cloning, reagents for restriction digest, PCR, and Gibson assembly were obtained from New England Biolabs, as was 5-α competent *Escherichia coli*. Oligos and gBlocks were obtained from Integrated DNA Technologies. ZymoPure Midiprep kit was used to prepare injection-quality plasmids and was obtained from Zymo Research. Plasmids were confirmed by Sanger sequencing and utilized generally available methods. We provide annotated sequences of the donor plasmids and genomic target regions in ApE format^31^ (at https://github.com/jchamper/ChamperLab/tree/main/Distant-Rescue).

### Generation of transgenic lines

Embryo injections were provided by Rainbow Transgenic Flies. The donor plasmid was injected into *w*^*1118*^ flies along with two helper plasmids, one providing gRNAs driven by the U6:3 promoter and one providing Cas9 driven by the *hsp70* promoter (plasmid sequences for these can also be found at https://github.com/jchamper/ChamperLab/tree/main/Distant-Rescue). Flies were housed with Cornell Standard medium in a 25°C incubator on a 14/10-hour day/night cycle.

### Genotypes and phenotypes

Flies were anesthetized with CO_2_ and screened for fluorescence using the NIGHTSEA adapter SFA-GR for DsRed and SFA-RB-GO for EGFP and ECFP (it was found to provide better results than SFA-VI). Fluorescent proteins were driven by the 3xP3 promoter visualization in the white eyes of *w*^*1118*^ flies. DsRed was used as a marker to indicate the presence of the drive alleles, and EGFP was used to indicate the presence of the supporting *nanos*-Cas9 allele^32^. ECFP was used for the *Rpl35A* rescue-only element.

### Phenotype data analysis

To calculate several statistics and rates in our study, offspring from different vials were pooled together before analysis. However, this approach did not take potential batch effects into account (different vials could be considered different batches of progeny), which could bias rate and error estimates. To account for this, we conducted an alternate analysis as in the previous studies^2,15,16,33,34^. We fit a generalized linear mixed-effects model with a binomial distribution (by maximum likelihood, Adaptive Gauss-Hermite Quadrature, nAGQ = 25). This model allows for variance between batches, resulting in slightly different parameter estimates and slightly increased standard error estimates in most cases (any large batch effects would tend to increase these further). This analysis was performed using the R statistical computing environment (3.6.1), including packages lme4 (1.1-21, https://cran.r-project.org/web/packages/lme4/index.html) and emmeans (1.4.2, https://cran.r-project.org/web/packages/emmeans/index.html). The script is available on Github (https://github.com/jchamper/ChamperLab/tree/main/Distant-Rescue). The resulting rate and error estimates were similar to the pooled analysis (Data Sets S1-5).

### Genotyping

To genotype flies, they were frozen, and DNA was extracted by grinding in DNAzol according to the manufacturer’s protocol (ThermoFisher Scientific). The DNA was used as a template for PCR using Q5 Hot Start DNA Polymerase from New England Biolabs. The region of interest containing gRNA target sites in *RpL35A* was amplified using DNA oligo primers RpL35ALeft_S_F and RpL35ARight_S2_R. This would allow amplification of wild-type sequences and sequences with resistance alleles but would not amplify recoded drive sequences because the reverse primer is outside the rescue element of all TADE drives. For all constructs, primers were used that would only produce PCR products if the genomic insertion was located where it was expected (see Supplemental Information).

## Results

### Drive constructs

Several transgenic constructs were used in this study. One was a split homing drive targeting *RpL35A* with rescue and has been described previously^2^. Another was a same-site TARE drive targeting *hairy*^16^. In this study, we generally used split drives that were coupled with a split *nanos*-Cas9 allele with an EGFP marker^32^ (Figure 1). A similar Cas9 allele was driven by the *vasa* promoter, though it retained the *nanos* 3’UTR.

**Figure 1.**
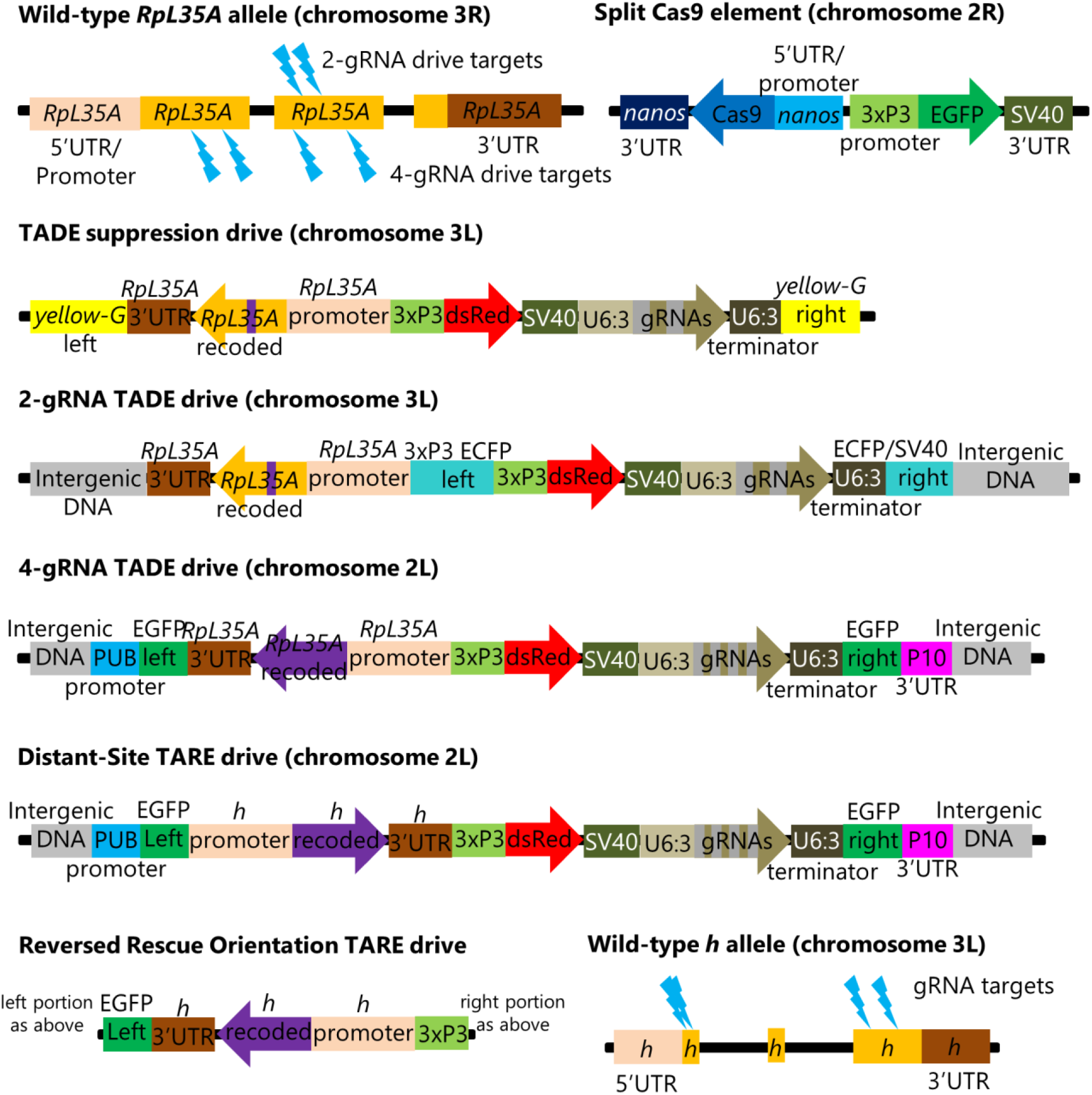
Genetic Constructs and target sites. The figure shows each element in several genetic constructs described in this study. Besides these, there is a *RpL35A* rescue-only construct that is similar to the 2-gRNA TADE drive but lacks the gRNAs and has an intact ECFP fluorescent protein gene instead of DsRed. Another construct is similar to *nanos*-Cas9, but with the *nanos* promoter and 5’UTR replaced by *vasa* elements.

Several distant-site TADE constructs were designed, injected, and confirmed to be inserted at the correct genomic site (see Supplemental Information). One consisted of a TADE suppression drive inserted at the *yellow-G* locus (Figure 1), which was previously used as a target for homing suppression drives^34,35^ because it is a haplosufficient but essential female fertility gene. This drive had the same pair of gRNAs as the homing drive targeting *RpL35A*. It had a full rescue element that included 486 nucleotides of DNA upstream of the first exon. The next closest gene (*POLDIP2*) started its 5’UTR 676 nucleotides upstream of the first *RpL35A* exon, so the promoter likely contained all *RpL35A*-specific regulatory elements. Only a small portion of the rescue element was recoded (a 39-nucleotide strength in between the two gRNA target sites).

A TADE modification drive was formed by first inserting a *RpL35A* rescue element as above (and with an ECFP fluorescent marker) at an intergenic site on chromosome 3L that previously supported effective homing at an EGFP target site^32^. A DsRed and gRNA set as above were then inserted inside the ECFP gene (Figure 1).

A second TADE modification drive was inserted into a polyubiquitin-EGFP gene at in intergenic region in chromosome 2L that had also supported a synthetic target homing drive^33^. However, this drive had four gRNAs that targeted different sites (Figure 1). The entire coding sequence was also recoded, and only the first two introns were retained (these introns separated exons containing the 5’UTR).

A distant-site TARE drive was inserted into the same spot as the 4-gRNA TADE drive. It also had four gRNAs, including two gRNAs used in earlier TARE drives^15,16^, an additional gRNA that was used for initial insertion of these previous same-site TARE drives^15,16^, and a final gRNA targeting the beginning of the coding sequence (Figure 1). The recoded rescue element was largely the same as the previous drives, but an additional 39 nucleotides of sequence was recoded after the start codon to prevent homology with the wild-type allele on both sides of the first gRNA cut site. 1718 nucleotides upstream of the 5’UTR (as measured from the isoform with the longest 5’UTR) were retained for use as possible promoter sequence for the rescue element.

A final TARE construct was as above, except that the orientation of the entire rescue element was reversed (Figure 1) to assess the potential impact of adjacent regulatory elements or transcription from the EGFP gene fragment.

### TADE suppression drive has low cleavage rates

To assess the drive efficiency, female TADE suppression drive carriers (which could be heterozygous or homozygous) were crossed to male Cas9 line homozygotes. Female and male offspring with both red and green fluorescence were selected and crossed to *w*^*1118*^ flies. They were allowed to lay eggs for one week and then removed. Progeny were collected and phenotyped. The drive inheritance rate, indicated by the proportion of offspring with DsRed, was 54% for all crosses that produced viable offspring (Figure 2A, Data Set S1), which was not significantly different than the Mendelian expectation (*p* = 0.1, Fisher’s exact test). This could indicate that the drive rescue element failed or that the toxin element of the drive did not function correctly (meaning that Cas9 cleavage did not occur).

**Figure 2.**
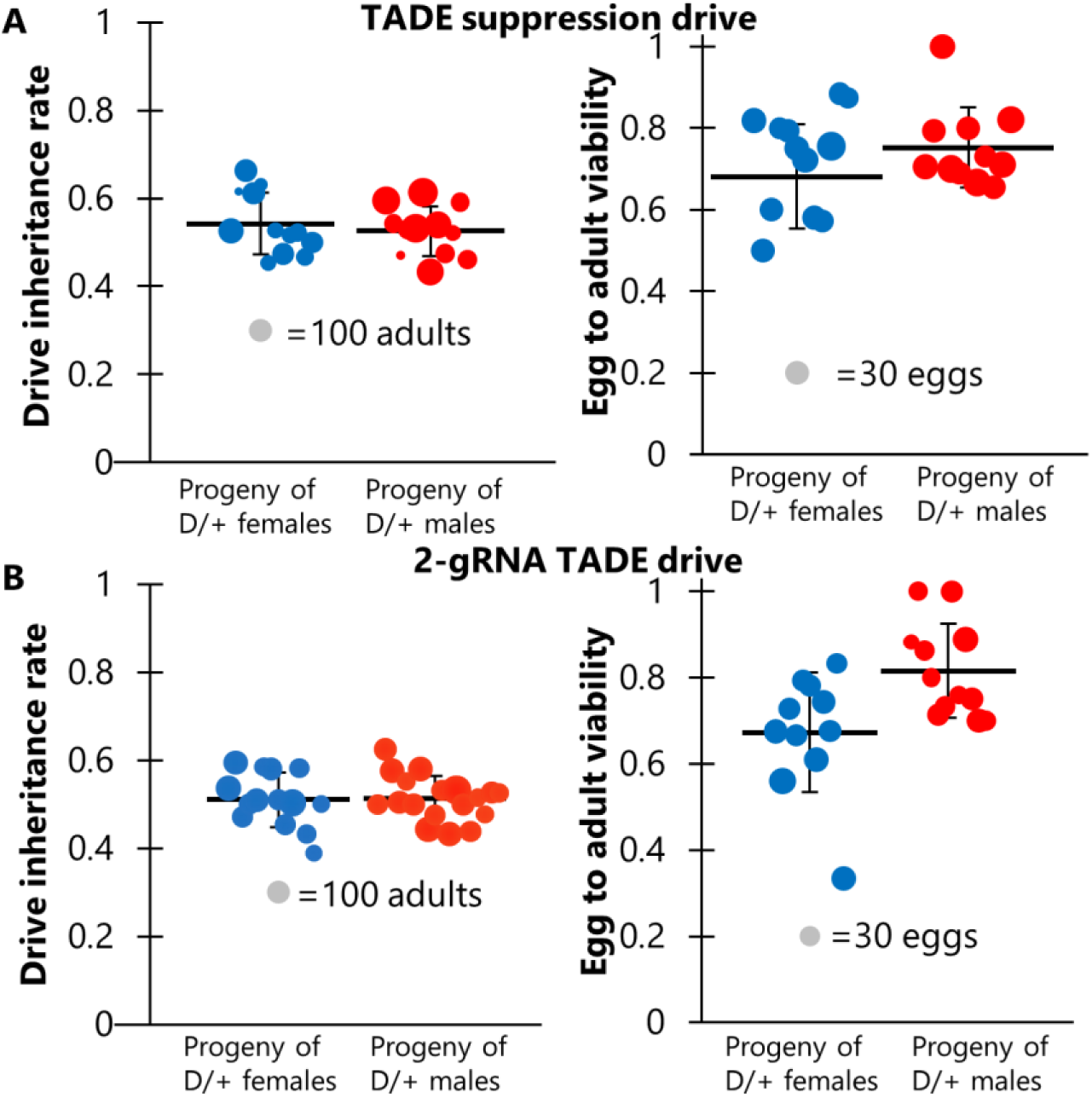
TADE drive inheritance and egg viability. Cross data are displayed for **(A)** the TADE suppression drive and **(B)** the 2-gRNA TADE drive. Drive inheritance and egg-to-adult viability was measured from crosses between drive individuals (heterozygous for the drive and for a Cas9 allele) and wild-type flies. Each dot represents offspring from one drive parent, and the size of dots is proportional to the number of total offspring for drive inheritance or the number of eggs for egg viability. Rate and standard error of the mean are displayed for all flies pooled together.

To assess the latter possibility, we examined egg viability in additional crosses of the same type, comparing them to crosses between *w*^*1118*^ flies. Flies were allowed to lay eggs for a period of 20-24 hours and were then transferred to a new vial for three consecutive days. After each transfer, the number of eggs was recorded, and all adults were later phenotyped. We found that egg viability was significantly lower in female and male drive crosses (72% and 75%) than in *w*^*1118*^ crosses (83%) (Figure 2A, Data Set S1). This could potentially be explained by a ∼10% germline cut rate and a ∼2% embryo cut rate (though the embryo cut rate is even more uncertain because female and male drive crosses did not have significantly different viability) in a model where any cut inherited by offspring results in nonviability. These results are consistent with target site sequencing of six progeny from male drive crosses, which found only wild-type alleles. These low rates make it unclear if the rescue element functioned effectively. Such cut rates are substantially lower than the 82% germline and 29% embryo rates of the homing rescue drive targeting the same *RpL35A* gene using the same two gRNAs and Cas9 allele^2^. This difference could perhaps be explained by the different genomic location and environment of the gRNA gene in this construct. Another possibility is that some cuts resulted in homology-directed repair using the wild-type allele as the template, with was not possible in the homing drive. This would mean that actual cut rates would be higher, but the effective disruption rate of the target site would remain low unless one of the alleles were already disrupted in the germline (and thus available as a template for homology-directed repair), though such genotypes could not be recovered for testing.

### Undesired homology-directed repair in a TADE modification drive

A simpler distant-site 2-gRNA TADE drive designed for modification was constructed (Figure 1), and we performed a similar set of crosses. Drive inheritance for these was 51-52%, close to the Mendelian expectation (Figure 2B, Data Set S2). However, the viability of eggs from drive females (66%) was significantly reduced compared to male egg viability (81%, p < 0.0001, Fisher’s exact test) and viability in *w*^*1118*^ crosses (83%, p < 0.0001, Fisher’s exact test) (Figure 2B, Data Set S2).

This is consistent with a 10% embryo resistance allele formation rate (in which a single resistance allele is sufficient to render the egg nonviable), or perhaps a 20% rate if only one cut site is available. However, germline rates have been generally higher than embryo cut rates in all studies, meaning that the lack of reduced viability in the progeny of males was unexplained. To assess this, we sequenced the target site of eight progeny from male drive crosses. All of these had the recoded sequence of the rescue element between the two cut sites. This means that the target on chromosome 3R was able to use the drive on chromosome 3L as a template for homology-directed repair (for which at least several hundred nucleotides would be available on either side of the cut sites), perhaps even from the same chromosome molecule. Despite apparently fairly high cut rates, this explains the full viability of progeny from male drive crosses and the relatively high viability of progeny from female drive crosses.

### TADE modification drive with high cleavage rates shows failure of distant-site rescue elements

To solve the issue of low-cut rates and the use of the rescue element as a template for homology-directed repair, we constructed a distant-site 4-gRNA TADE modification drive (Figure 1). This drive has a much larger recoded region and used four completely new gRNAs. When drive and Cas9 heterozygotes were crossed to *w*^*1118*^, no viable progeny were obtained for both male and female crosses, despite several eggs being laid (Figure 3A, Data Set S3). This indicates that the drive has a 100% rate of germline disruption, much higher than our TADE suppression drive. However, it also indicates that the drive was unable to provide rescue.

**Figure 3.**
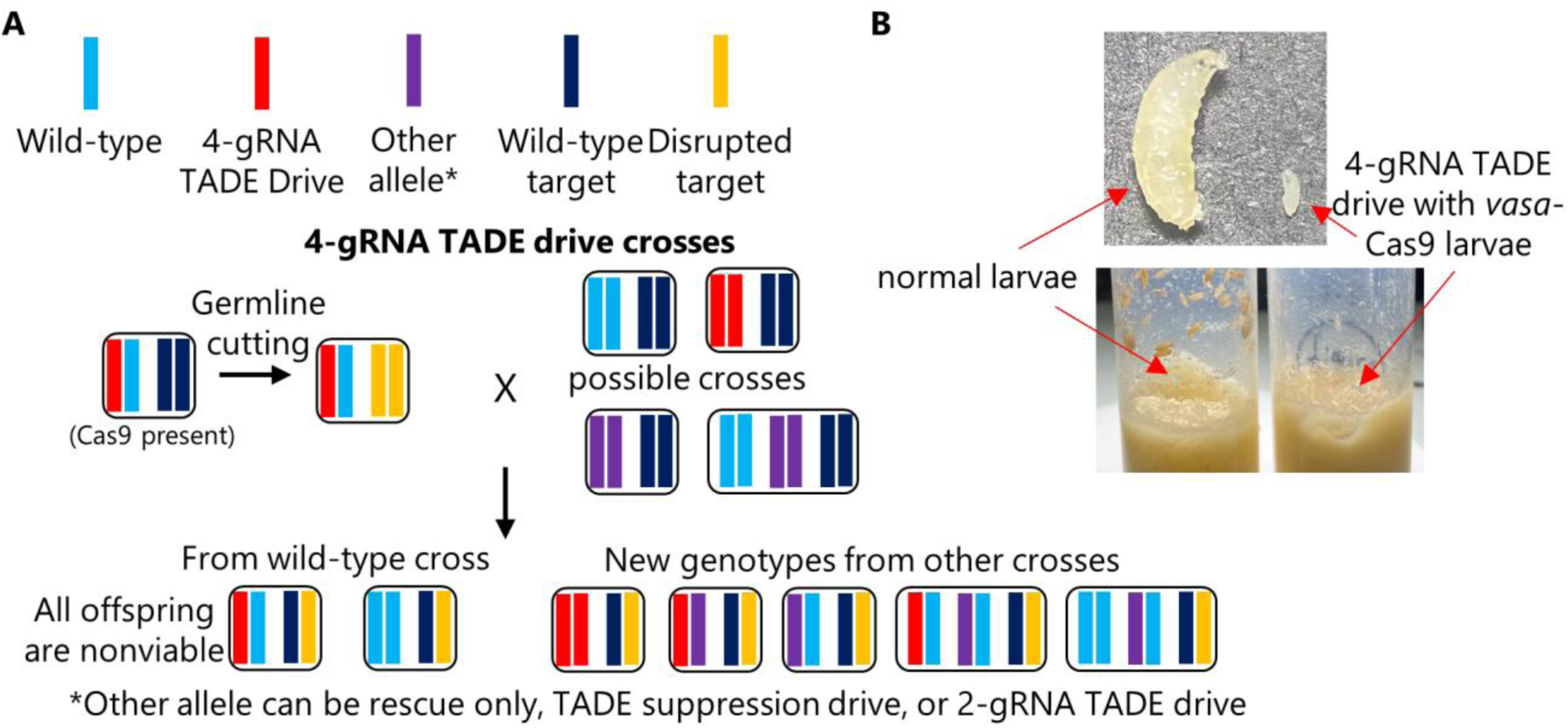
TADE rescue assessment. (**A**) Individuals heterozygous for the 4-gRNA TADE drive and the *nanos*-Cas9 allele were crossed to flies of several genotypes, including *w*^*1118*^ and three genotypes that could provide different distant-site rescue elements for the *RpL35A* target gene. However, no progeny were viable, indicating that all rescue elements failed. (**B**) When flies homozygous for the 4-gRNA TADE drive were crossed to flies homozygous for a *vasa*-Cas9 allele, offspring suffered from leaky somatic expression and failed to develop, with larvae growing more slowly and never reaching significant size (the maximum size reached is shown in the image).

An additional cross between drive/Cas9 males and drive homozygous females also failed to yield progeny, indicating the drive was unable to provide partial rescue. We also obtained no progeny from similar crosses between drive/Cas9 males and female homozygotes for the rescue-only allele, females homozygous for the TADE suppression drive, and females homozygous for the 2-gRNA TADE modification drive (Figure 3A, Data Set S3). Thus, none of our constructs had a functioning rescue element. We also obtained no progeny when these males were crossed to proven homing rescue drive females, though this was expected because the successful same-site rescue element could likely not express *RpL35A* more efficiently than the wild-type allele.

### Alternate Cas9 promoters and somatic expression

Because TADE drive targets a haplolethal gene, even a moderate amount of somatic expression may be sufficient to prevent success of the drive. To assess this, we generated a Cas9 allele that was nearly identical to the *nanos*-Cas9 construct but utilized the *vasa* promoter and 5’UTR elements in place of *nanos* upstream elements (the *nanos* 3’UTR was retained). In flies, *vasa* is known to be a strong germline promoter but also has moderate somatic expression compared to *nanos*, which has little to no expression in somatic cells^36,37^.

To investigate the impact of somatic expression on the 4-gRNA TADE drive, drive homozygous females were crossed to males that were homozygous for the *vasa*-Cas9 allele, and they were allowed to lay eggs in several vials. All offspring thus possessed both drive and Cas9 alleles. We found that eggs usually survived and hatched into larvae. However, all larvae had growth defects and never reached a substantial size (Figure 3B) before eventually expiring several days after they would normally have become pupae. This indicates that any TADE drive (or at least those with a strong toxin element) should use promoters that have little to no somatic expression.

### Distant-site TARE drive assessment

Though haplolethal genes require precisely controlled expression to achieve high fitness, haplosufficient genes can tolerate a wider range of expression. Thus, we tested a distant-site TARE drive to see if distant-site rescue might function correctly.

This drive used the same target gene and promoter region as previous TARE drives^15,16^ but used two additional gRNAs for a total of four. As expected, crosses between *w*^*1118*^ females and males with one copy of the drive and one copy of Cas9 had normal numbers of offspring and no biased inheritance (Figure 4A, Data Set S4). Such progeny would still possess one wild-type *hairy* allele from the female parent and thus be viable. However, crosses between *w*^*1118*^ males and females with one copy of the drive and one copy of Cas9 produced no offspring. In these crosses, high rates of maternal Cas9 and gRNA deposition resulted in complete or nearly complete cleavage of wild-type *hairy* alleles in all progeny, rendering them nonviable. We can conclude that the rescue element failed because a functional rescue element would have allowed drive-carrying offspring to survive. Even when these drive/Cas9 females were crossed to drive homozygote males, no viable progeny were obtained (Data Set S4), indicating that the distant-site TARE drive does not even provide partial rescue. In contrast, when crossed to previous functional TARE drive homozygotes^16^, all progeny were viable (one gRNA of the new drive may have been able to cleave the old TARE drive allele, but must have done so at a low rate in the embryo).

**Figure 4.**
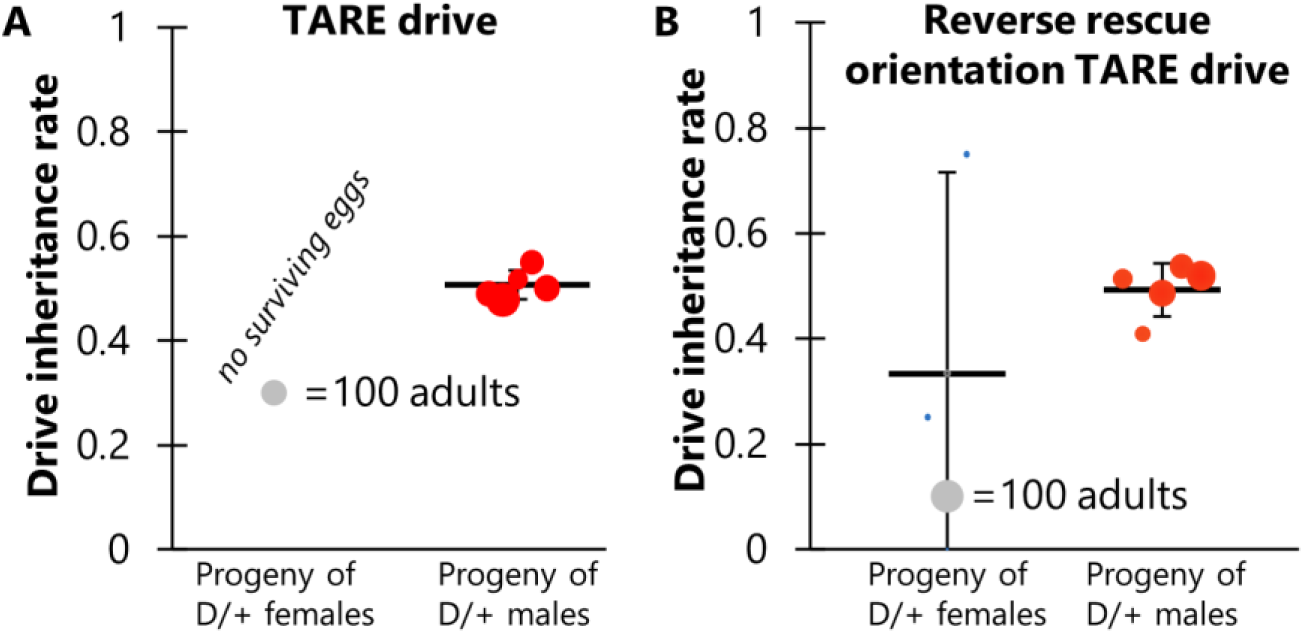
TARE drive inheritance. Drive inheritance was measured from crosses between drive individuals (heterozygous for the drive and for a Cas9 allele) and wild-type flies. Each dot represents offspring from one drive parent, and the size of dots is proportional to the number of total offspring. Rate and standard error of the mean are displayed for all flies pooled together

Because the polyubiquitin promoter element and EGFP fragment may have interfered with rescue element expression in the TARE drive, we constructed an identical allele in which the orientation of the rescue element was reversed (Figure 1). However, crosses involved this construct showed nearly identical results to the first TARE drive, except that a few female drive cross vials had very small numbers of surviving offspring, averaging less than one per vial (Figure 4B, Data Set S5). Thus, the rescue element could not function in reverse orientation.

Of note, both of these 4-gRNA TARE drives showed ∼100% germline cutting and 98-100% embryo cutting rates, as indicated by the lack of surviving offspring (Data Set S4-5 - this estimate is based on the requirement of cutting both wild-type alleles in progeny to render the progeny nonviable, as well as many previous observations that show higher germline cut rates than embryo cut rates). This represents a substantial increase from the estimated 89% germline and 63% embryo cutting of the complete TARE drive^16^, or the ∼100% germline cutting and ∼86% embryo cutting of the split TARE drive that used the same Cas9 element^15^. This further indicates that additional gRNAs can substantially increase total cut rates in systems where weak gRNAs may be a limiting factor.

## Discussion

In this study, we examined the possibility of developing distant-site TADE suppression and modification drives targeting *RpL35A* and TARE drives targeting *hairy*. Though improved gRNA cassettes increased cut rates compared to previous drives targeting these genes, the drives could not function because their rescue elements were not effective. Additionally, the small recoded region in one drive could be used as a template for homology directed repair of the cleaved wild-type target allele, thus forming a functional resistance allele.

The need for effective rescue is absolutely essential for construction of any toxin-antidote drive, and the ability to perform this rescue from a site distal from the target gene would increase potential options. For example, it would preclude the need for additional gRNAs targeting the female fertility gene in TADE suppression drive^19,20,25^, which would also reduce the effects of leaky Cas9 somatic expression on female fertility^38^ (though such effects may be small compared to leaky somatic expression on the haplolethal TADE target, as demonstrated by our use of the *vasa* promoter). It would also enable to construction of 1-locus 2-drive systems, which are highly confined^20^. However, none of our systems was able to provide effective rescue. Distant site rescue is certainly possible, as demonstrated for several haplosufficient but essential genes in ClvR drive systems^17,18^. All of these were located at the same genomic site, so it is possible that certain genomic sites are more amenable to correct rescue expression than the ones we tested. Our own TARE drive with a haplosufficient target was only tested at one site. This site was known to support previous successful homing drives^33^, but it is possible that the polyubiquitin promoter interfered with rescue expression (despite testing the rescue element in both possible orientations). Thus, even though our test failed, distant-site rescue for haplosufficient target should still be considered a viable option. Possible improvements could include longer promoter elements, other stronger rescue expression systems, new target genes, better genomic insertion sites for the drive, or flanking of the rescue element with sequences to protect it from local genomic structure (such as gypsy elements).

For haplolethal targets, precise expression is even more important, potentially increasing the difficulty of successful rescue. One of our TADE rescue elements was similar to the TARE element and may have failed for the same reasons. However, our other two rescue elements were at very different sites, one site a native gene and both known to support effective homing drives^32,34^. Neither were near the polyubiquitin promoter, and both had minimal recoding, meaning that potentially important introns and other sequences were preserved. Yet, both of these failed, which suggests that distant-site rescue elements for haplolethal genes may be considerably harder to construct. In this case, they likely failed through underexpression rather than overexpression, because having either one or two additional copies of the rescue element could not provide rescue for even one disrupted wild-type allele, nor did drive homozygotes show any phenotype associated with negative fitness effects when they were homozygous for wild-type *RpL35A*. Further underscoring the difficulty of successful rescue was our observation of homology-directed repair between the rescue element of one of the drives with minimal recoding on chromosome 3L and the target site on chromosome 3R. This indicates that recoded regions, even at distant sites (and perhaps even in different chromosomes, considering the distance between arms in chromosome 3) should probably be highly recoded, with as many mutations as possible on both sides of the target site for at least a few hundred nucleotides for any CRISPR/Cas9 toxin-antidote drives, thus minimizing the chance of using the drive’s rescue element as a template for homology-directed repair. However, this may not be the case for all genomic sites because we did not observe this phenomenon in our TADE suppression drive.

Several of our previous gene drives in *Drosophila* with the same *nanos*-Cas9 promoter had nearly 100% germline cut rates and generally high embryo cut rates when targeting the *yellow*^32,39,40^, *white*^36^, *cinnabar*^36^, and synthetic EGFP^32,33^ genes. Yet, more recent successful drives targeting *hairy*^15,16^ and *RpL35A*^2^ had cut rates that were considerably below 100% in the germline and embryo, despite having two gRNAs. Our TADE drives using the same two 2-gRNA cassettes demonstrated further variation, with one having high germline cutting based on sequencing results and the other having fairly low cutting rates. This indicates that gRNAs, in addition to Cas9^32,33^, can have varied expression based on their genomic location. For both our TARE and TADE drives, replacing the 2-gRNA cassette with a 4-gRNA cassette (and retaining the same U6:3 promoter and tRNA separation system) aiming at the same region of the target gene yielded nearly 100% cut rates. Strictly speaking, this cutting increase can only be confirmed to take place in the germline of the TADE drive and the early embryo of the TARE drive, but it was generally a successful strategy. Even if additional gRNAs are not necessarily needed to avoid functional resistance alleles, they can provide insurance against low-activity gRNAs without any significant downsides for CRISPR toxin-antidote drives^19^ (unlike homing drives^33^). However, caution is advised in this approach for TADE drives because if the embryo cut rate increases (as it did for our TARE drive), the confinement level of the drive will also be considerably increased^19,20,25^, which may not be desired.

In conclusions, we found that developing efficient distant-site rescue elements may be more challenging than anticipated, particularly for TADE drives. This may necessitate trying multiple sites for TARE drives using methods such as PiggyBac transformation and focusing efforts on developing same-site TADE suppression drives rather than potentially simpler distant-site systems. To achieve high germline cut rates, additional gRNAs could be added to toxin-antidote drives beyond the amount needed for functional resistance allele avoidance.

## Acknowledgements

This study was supported by laboratory startup funds from Peking University and the NSFC Overseas Youth Fund.

## Supplemental Information

### Plasmid key and injections

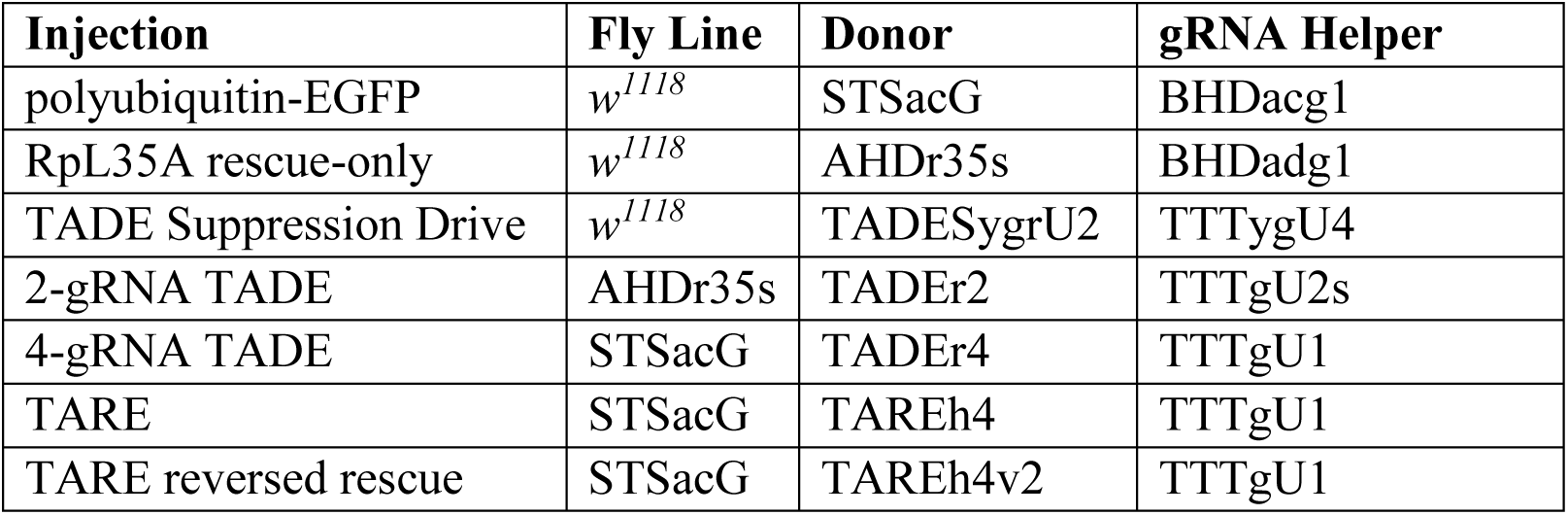

### *Rpl35A* target site analysis primers

~~~
RpL35ALeft_S_F: GCATGCAAATGATCGAAACCCT
RpL35ARight_S2_R: CGTTTCCATCGTCTTCATCTGC
~~~

### STSacG insertion site check primers

~~~
AutoCLeft_S2_F: AGATTGGCCACCACATCCATC
EGFPaLeft_S_R: GCTTGTTTATTTGCTTAGCTTTCGC
~~~

### AHDr35s insertion site check primers

~~~
AutoDLeft_S2_F: TTTGATCTGATTGCGACGCGT
RpL35ARight_S_F: CGCATCCGCATCGTTAGTTCA
~~~

### TADESygrU2 insertion site check primers

~~~
U6term_S_F: CATCTGACGTGTGTTTATTTAGAC
YGRight_S2_R: TAATGAGACCCAGTAACGACA
~~~

### TADEr2 insertion site check primers

U6term_S_F: CATCTGACGTGTGTTTATTTAGAC

AutoDRight_S2_R: GCAATGGTAATGACTCACAGT

### TADEr4, TAREh4, and TAREh4v2 insertion site check primers

~~~
U6term_S_F: CATCTGACGTGTGTTTATTTAGAC
AutoCRight_S_R: TACACCTCACACTACTCGGGC
~~~

## Notes

https://github.com/jchamper/ChamperLab/tree/main/Distant-Rescue

### Competing Interest Statement

The authors have declared no competing interest.

